# The Effect of Between-Probe Variability in Haptic Feedback on Stiffness Perception and Grip Force Adjustment

**DOI:** 10.1101/2020.06.15.151605

**Authors:** Hanna Kossowsky, Mor Farajian, Amit Milstein, Ilana Nisky

**Affiliations:** Department of Biomedical Engineering, Ben-Gurion University of the Negev, Beer-Sheva, Israel

**Keywords:** Computational models, grip force, stiffness perception, uncertainty, variability

## Abstract

When interacting with objects, haptic information is used to create perception of the object stiffness and to regulate grip force. Studies have shown that introducing noise into sensory inputs can create uncertainty in those sensory channels, yet a method of creating haptic uncertainty without distorting the haptic information has yet to be discovered. Toward this end, we investigated the effect of between-probe haptic variability on stiffness perception and grip force control. In a stiffness discrimination task, we added different levels of between-probe haptic variability by changing the stiffness of the force fields between consecutive probes. Unlike the low and high variability levels, the medium level created perceptual haptic uncertainty. Additionally, we ascertained that participants calculated a weighted average of the different stiffness levels applied by a given force field. Examining participants’ grip force showed that the modulation of the grip force with the load force decreased with repeated exposure to the force field, whereas no change in the baseline was observed. These results were observed in all the variability levels and suggest that between-probe variability created haptic uncertainty that affected the grip force control. Overall, the medium variability level can be effective in inducing uncertainty in both perception and action.

## I. Introduction

When interacting with objects in our surroundings, we receive feedback from multiple senses and integrate them to create perception and action. Haptic sensory information is used to estimate external forces acting on our limbs and the mechanical properties of objects [1-3]. An example of such a mechanical property is the stiffness of elastic objects, i.e., the linear relationship between the penetration into the object and the resulting force. To estimate the stiffness of an object, our brain integrates force and position information obtained from the sensory system [1, 4-7]. Based on these estimates, we create our perception of the stiffness of an object. These estimates are also used by our motor system to regulate actions, such as grip force, which is the force that we apply perpendicularly between our fingers and an object to maintain a stable grasp.

Previous studies have utilized programmable robotic devices to investigate how our haptic perceptions and actions are created [8]. Using these devices to present participants with systematically manipulated stimuli, researchers have investigated the human motor control system by recording participants’ hand positions and applied forces. The formation of perception can be studied by logging participants’ answers to perceptual judgment questions [9, 10].

Presenting participants with altered sensory inputs has enabled researchers to study the ways in which the nervous system forms perception. It is possible to identify the computational model that is most likely to be in accordance with the neural processes by positing various computational models, and isolating the model which best fits participants’ perceptual estimations [5-7]. For example, various models have been suggested for stiffness estimation when a delay was introduced between force and position information [5, 7].

Kuschel et al. [6] modified the relation between visual and haptic information, showing that visual-haptic integration in compliance estimation is likely a weighted sum of the visual and haptic sensory inputs. In a study by Metzger et al. [11], the researchers manipulated the stiffness of a single probe out of a series of identical multiple probes carried out by each participant. This allowed them to determine that there are cases in which the nervous system integrates serial haptic information as a weighted average. Their model attributes decreasing weights to the different stiffness levels based on their serial placement in the probing sequence, with the highest weight given to the stiffness level experienced first and the lowest to the last. With the exception of Metzger et al. [11], the integration of haptic information received through multiple interactions with the same object has received little attention in the context of stiffness perception.

One prevalent method used to investigate perception and action is to introduce uncertainty into the sensory inputs [12]. The effect of this induced uncertainty on the perception of object properties and on actions can shed light on the processes by which they are formed. The creation of uncertainty can be achieved by adding noise to sensory inputs [12, 13]. Such artificial noise is added to the inherent noise in the human motor and sensory systems [14]. By adding varying levels of artificial noise to visual information, Ernst and Banks [12] studied how visual and haptic information are integrated in height perception. Perception is a probabilistic process, and hence, the estimate of any property will have an associated variance. They showed that the integration of visual and haptic information is a convex combination that weights each sense proportionally to its reliability, such that the integration is statistically optimal. Gurari et al. [13] found that adding noise to force and stiffness haptic stimuli led to a degradation in the ability to perceive these stimuli.

We propose that haptic uncertainty can also be created by varying the mechanical properties of an object between consecutive interactions rather than by adding noise within a given interaction. That is, we suggest preserving the haptic information presented in each probing movement (i.e., no noise is added) and investigating the possibility of creating haptic uncertainty by varying the undistorted information between consecutive probing movements. If this method is found to create uncertainty in haptic interactions, it may open the possibility of investigating haptic information processing without necessitating qualitative changes in the haptic information that is presented during an individual probing movement. To the best of our knowledge, previous studies have not investigated how between-probe variability in haptic information affects perception and action. In this work, we make use of this method and investigate the effect of this variability on both the perception of stiffness and the control of grip force.

During object manipulation, we apply grip force to prevent the object from slipping due to load forces (e.g., gravity forces [15-19], environment interaction forces [20-23], and inertial forces [17, 24]) which act on the object. The grip force is mostly predictive [25, 26] and comprised of two components: (1) a baseline and (2) a modulation of the grip force in anticipation of the load force [22, 27, 28]. The baseline component is the grip force applied when there is no load force acting. The modulation component combines the internal representation of the object with that of the environmental dynamics to predict the load forces that may act on a manipulated object. This internal representation is formed by the nervous system [25, 29-31] and is updated during repeated interactions with an object [27, 28, 32].

Our grip force is adjusted in anticipation of the load force applied on our hand by an object [20, 22, 28, 33-36]. The load force depends on the mechanical properties of the object such as its mass and stiffness [9, 37]. Flanagan et al. [10, 38] demonstrated a tight coupling between grip force and load force for movements in different directions and rates, and showed that the maximum grip force and maximum load force coincide. Additionally, the friction between the finger pads and the object affects the amount of grip force that is necessary to prevent its slippage [9, 39-41]. Hence, real and virtual changes in these mechanical properties have been shown to affect grip force control [9, 22, 33, 42].

Grip force is additionally affected by one’s certainty regarding the object’s mechanical properties. The measure of experienced uncertainty defines the safety margin, which is difference between the applied grip force and the minimal grip force necessary to prevent the slippage of the object [9]. An increased uncertainty regarding the load force dynamics can lead to an increase in the safety margin [42]. This increase would come across in both the baseline and the modulation components. Although the baseline component does not depend directly on the anticipated load force, it is affected by the uncertainty regarding the object’s mechanical properties [42, 43]. The modulation component is affected by both the mechanical properties of objects and the uncertainty concerning the internal representation of these mechanical properties. Hence, in the event of uncertainty about the load force, both grip force components, the baseline and the modulation, would be affected.

Hadjiosif and Smith [42] investigated the adjustment of grip force to velocity dependent load forces that varied between subsequent reaching movements in a force field adaptation study. They showed that the grip force is adjusted based on both the expected load force dynamics and the variability of the load force. Furthermore, they found grip force to be threefold more sensitive to the variability of the load force, than to its expected value. To the best of our knowledge, the effect of varying elastic force fields on grip force is yet unknown.

In this work we aim to develop a method of introducing uncertainty into haptic information without distorting the haptic feedback supplied in each individual probing movement. We do this by varying the haptic feedback between consecutive probes into a given force field, such that no noise is added to each separate probe. We conduct a stiffness discrimination experiment using a virtual reality setup and test the effectiveness of between-probe haptic variability as a method of inducing uncertainty into the participants’ perceptual estimations and their grip force adjustment. We hypothesize that higher levels of variability will lead to increased uncertainty, and suggest that this uncertainty will come across in both the stiffness perception and in the two different components of grip force control. The perceptual uncertainty may be expressed as a decrease in the discrimination sensitivity. Additionally, as the between-probe variability may increase participants’ uncertainty regarding the load force, we expect to find an increase in the grip force baseline to expand the safety margin. Furthermore, we expect to find a weaker or unchanged modulation of the grip force with the load force after repeated interactions with a force field whose stiffness level varies between consecutive probes. In addition, we test several simple models to investigate how the varied haptic information experienced during serial probing movements is combined to create the perceived stiffness.

## II. Methods

### A. Participants

Ten right-handed participants (N=10, 7 females, ages 22-29) completed the experiment after signing an informed consent form. The form and the experimental protocol were approved by the Human Subject Research Committee of Ben Gurion University (BGU) of the Negev, Be’er Sheva, Israel. The participants, students at BGU, were compensated for their participation in the experiment, regardless of their success or completion of the experiment.

### B. Experimental Setup

The aim of this study was to understand the effect of between-probe haptic variability on the perception of stiffness and on grip force control. Our experimental setup consisted of a custom-developed simulation in a virtual reality (VR) environment, using the Open Haptics API, written in C++ (Visual Studio 2010, Microsoft). Participants interacted with the virtual environment using a PHANTOM® Premium 1.5 haptic device (3D SYSTEMS, South Carolina, USA) and viewed the virtual environment through a semi-silvered mirror that blocked their view of their hand and showed the projection of an LCD screen placed above it, as depicted in Fig. 1. Throughout the experiment the participants wore noise cancelling headphones to eliminate auditory cues.

**Fig. 1.**
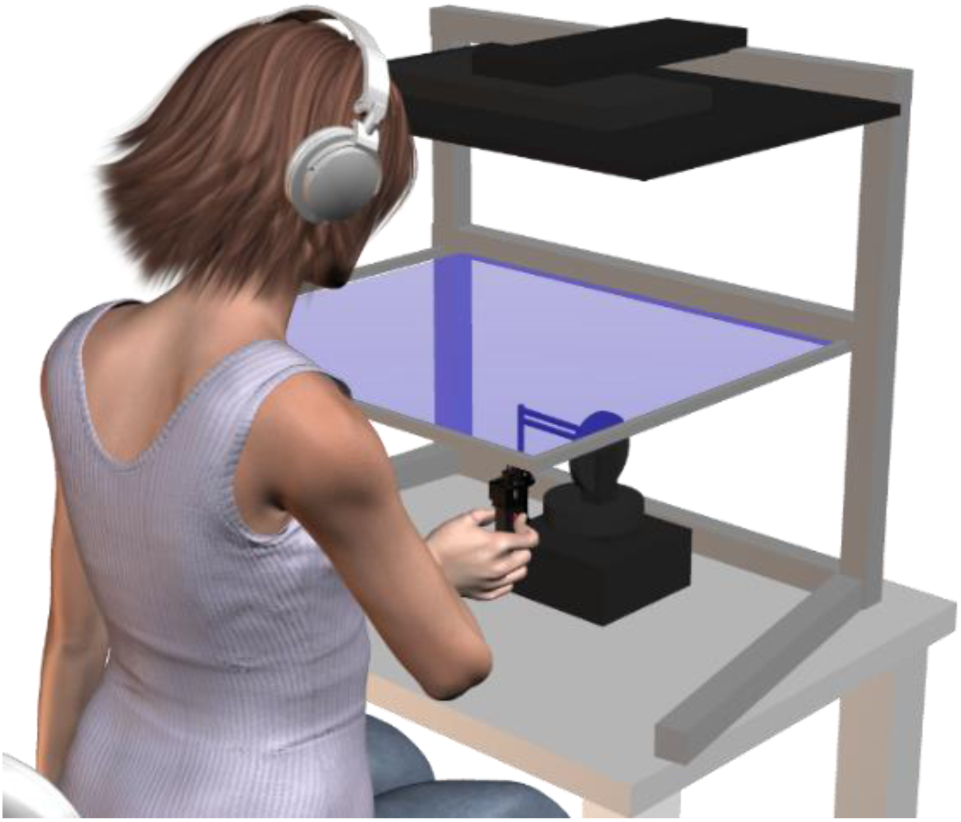
Experimental setup. The participants sat in front of a virtual reality system and used their right hand to grasp a haptic device and probe virtual elastic force fields. The LCD screen was either blue or red, to enable participants to differentiate between the two different force fields (standard and comparison) presented in each trial. Participants wore noise cancelling headphones to eliminate auditory cues. The force sensor was embedded in the black capsule, to allow the measurement of the applied grip force. We will note that the black capsule is the lower part of a device that can be used to create artificial skin stretch via moving tactors. This device was used in other studies with the same setup to study the dissociable role of the two haptic modalities (tactile and kinesthetic) [27]. In this work, participants grasped the device directly above the force sensor and not above the tactors, and the skin stretch stimulation was never actuated.

Participants held the haptic device in their right hand, using their index finger and thumb. The haptic device generated a virtual elastic force field that emulated a one-sided spring. The force field was rendered in the upward direction and was proportional to the distance between the tip of the haptic device and the virtual boundary of the force field. The haptic information was rendered at 1 KHz and was applied only when participants were in contact with the force field, which was defined to be the negative half of the vertical axis:

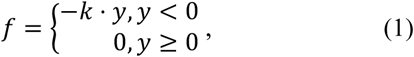

where k [N/m] is the stiffness, and y [m] is the penetration distance into the virtual force field. A force sensor (ATI, Nano 17), embedded in a plastic capsule mounted on the end of the haptic device, enabled the measurement of the grip force that participants applied between their thumbs and index fingers.

### C. Protocol

This experiment was a forced-choice paradigm stiffness discrimination task in which participants were asked to make downward vertical probing movements into a pair of virtual elastic force fields. The force fields were designated *standard* and *comparison*, and after interacting with each field twice, participants were asked to choose which of the two felt stiffer. Each of the force fields was indicated to the participants by the color of the LCD screen, which was pre-defined pseudo-randomly to be either red or blue (Fig. 1). The order of the force field (*standard* or *comparison*) was pseudo-randomly chosen prior to the experiment.

The participants performed a total of eight discrete probing movements in each of the two force fields in each trial. Only successful probes were counted. These were defined as probing movements that started and ended outside the elastic force field, were completed within 300 ms, and extended at least 20 mm into the force field. To avoid overheating of the robotic device, we used an auditory alert if the penetration into the force field exceeded 40 mm. After four successful probing movements into the first force field, the field automatically switched to the second force field for four probing movements. Following this, participants once again probed the first, and then the second, force field an additional four times. Thereafter, the participants reported which field had a higher level of stiffness by pressing a keyboard key with the color corresponding to the LCD screen color of the stiffer force field. Lastly, the participants began the next trial by raising the end of the robotic device to at least three cm above the boundary of the force field.

In each trial, the *comparison* field’s stiffness level remained constant throughout the trial and was selected to be one of eight values evenly spaced between 40-130 N/m. The *standard* force field stiffness in each trial was comprised of eight values drawn from a normal distribution with a mean of 85 N/m and one of four possible standard deviations [zero, low (13 N/m), medium (26 N/m) or high (39 N/m)]. The resulting *standard* stiffness varied between each of the probing movements within the trial (excluding the zero standard deviation trials) [Fig. 2]. As the participants thought they were interacting with the same object in each of the eight probing movements, while in reality the stiffness level varied between the probing movements, we hypothesized that this would increase the uncertainty regarding the haptic information.

**Fig. 2.**
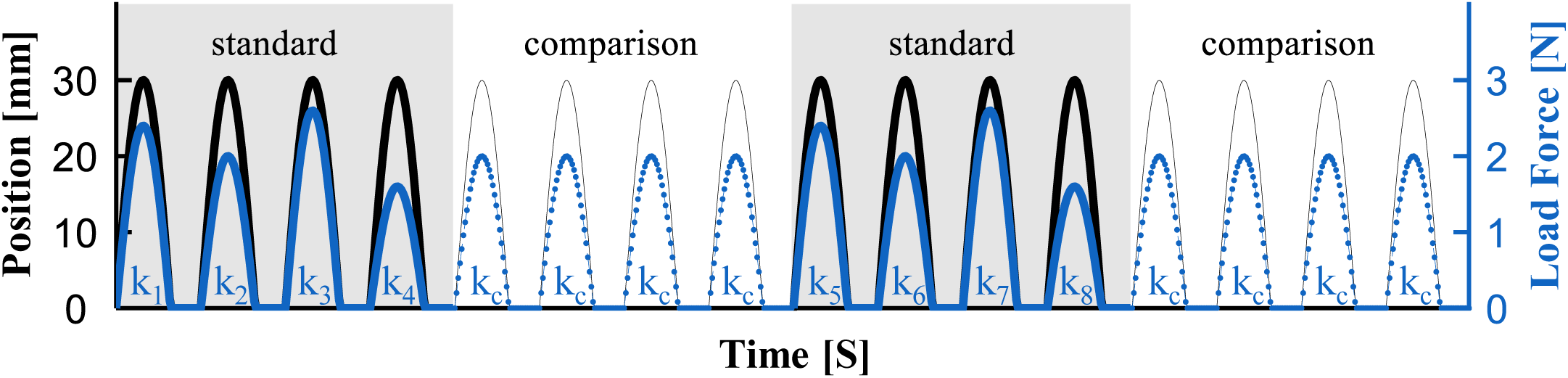
Illustrative position and load force trajectories for the two force fields. The black line represents the y position of the hand, and the blue line represents the load force that was applied by the haptic device. The gray shaded regions highlight the probing movements into the standard force field in which the stiffness level varied between consecutive probing movements throughout the trial. This caused different amounts of force to be applied for a given penetration distance. In the comparison force field (white regions), the stiffness (and hence the force that was applied for a given penetration distance) remained constant throughout the trial.

Previous studies have shown that the Just Noticeable Difference (JND) in stiffness perception, i.e., the sensitivity of humans to small differences between two stiffness levels, is approximately 15% of the stimulus value [44]. Based on this, we defined JND_L_=15% and the low, medium and high standard deviations corresponded to 1.JND_L_ (*σ* = 13 *N*/*m*), 2.JND_L_ (*σ* = 26 *N*/*m*) and 3.JND_L_ (*σ* = 39 *N*/*m*) of the mean stiffness level (85 N/m), respectively.

In total, there were eight *comparison* stiffness levels and four *standard* variability conditions, amounting to a total of 32 *standard*-*comparison* combinations. Each combination was repeated eight times throughout the experiment, leading to a total of 256 test trials, which were completed over two sessions of 144 trials each. The order in which the force field combinations were applied was pseudo-randomly chosen prior to the experiment. Half of the participants (N=5) completed the trials with the zero and medium standard deviations on the first day, and those with the low and high standard deviations on the second day, while the other half (N=5) completed the two sessions in the opposite order. Additionally, the participants began each session with 16 training trials, allowing them to become familiarized with the experimental setup and the haptic device. The training trials consisted of two repetitions of the eight *comparison* stiffness levels, and a zero standard deviation for the *standard* force field (i.e., a constant stiffness level of 85 N/m) and were excluded from the analyses. In the training trials participants received feedback about their response (correct or incorrect), whereas in the test trials, they received no feedback.

### D. Data Analysis

#### 1. Perception and Uncertainty

For each of the ten participants, we used the Psignifit toolbox 2.5.6 [45] to fit psychometric curves to the probability of responding that the *comparison* force field felt stiffer than the *standard* as a function of the difference between the stiffness levels of the two force fields:

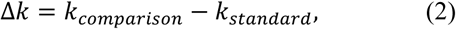

where *k*_*comparison*_ and *k*_*standard*_ are the *comparison* and *standard* stiffness values respectively. The probability was calculated as:

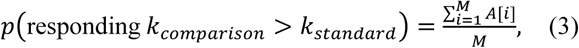

where M is the number of trials with a given Δ*k*, hence M=8 in this study. *A*[*i*] represents the participant’s answer for a given trial in a binary fashion:

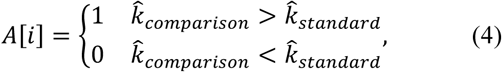

where 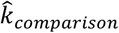 and 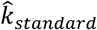 are the perceived *comparison* and *standard* stiffness values respectively. The resulting shape of the psychometric curve can be described as:

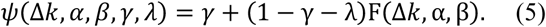

The shape of the psychometric curve depends both on the parameters *α, β, γ, λ* and on the function F, which is a logistic function in our case. We fit four psychometric curves to each of the participants’ responses, one for each of the four levels of variability.

Following this, we computed the Point of Subjective Equality (PSE) and the Just Noticeable Difference (JND) of each psychometric curve. The PSE is defined to be the stiffness level at which the probability of responding *comparison* is half, and therefore represents the measure of bias in the perceived stiffness:

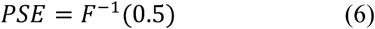

A rightward shift of a psychometric curve would indicate an increase, and a leftward shift – a decrease, in the perceived stiffness of the *standard* force field. The JND equals half the difference between the stiffness levels corresponding the 0.75 and 0.25 probabilities:

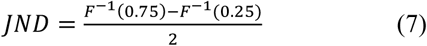

Hence, this value quantifies the slope of the psychometric curve, which represents the variability in participants’ responses, and is therefore an indication of the uncertainty experienced by the participants when choosing which force field was stiffer.

We examined the effect of the different levels of between-probe variability on the PSE and JND values across all the participants using a repeated-measures General Linear Model with the MATLAB statistic toolbox (2018a). The independent variables were the variability condition (categorical, df=3), and the participants (random, df=9). The dependent variables in the two separate analyses were the PSE and JND values. To compare the low, medium and high variability conditions to the zero-variability condition, we performed three planned t-tests using the Holm-Bonferroni correction for multiple comparisons. We present the p-values after this correction (*p*_*corrected*)_, and therefore the threshold significance level following the correction is 0.05.

#### 2. Models for Serial Stiffness Levels Integration

In order to confirm that the effect of the variability on the JND was due to participants interacting with a force field whose stiffness level was sampled from a normal distribution with a non-zero standard deviation, it was critical to ascertain if the participants used the information from most of the probing movements in a trial. Participants interacted with eight stiffness levels when probing the same *standard* virtual force field. We were interested in understanding whether their perceived stiffness was based on one stiffness level alone, e.g., the first or highest stiffness, or rather on all eight stiffness levels, e.g., an average of those values. To uncover the participants’ decision strategy, we developed computational models for different potential decision strategies, and ran simulations in which we compared the predicted answers of each of the models to those of each of the participants in each trial, with the goal of finding the models that best describe the participants’ perception.

Let us assume that for a given trial a participant interacted with the following sequence of stiffness levels during their interaction with the *standard* force field:

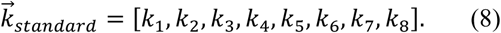

We considered the following nine computational models of perceived stiffness:

**Min** – the minimal of the eight stiffness levels:

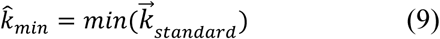

**Max** – the maximal of the eight stiffness levels:

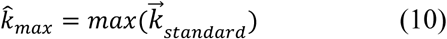

**Last** – the last of the eight stiffness levels:

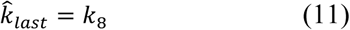

**First** – the first of the eight stiffness levels:

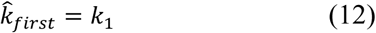

**Average of the Last Two** – the average of the last two of the eight stiffness levels:

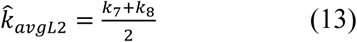

**Average of the First two** – the average of the first two of the eight stiffness levels:

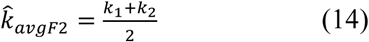

**Time Weighted Average** – we calculated the relative time that each of the eight stiffness levels was applied on each of the participants and used these values as weights in a weighted average:

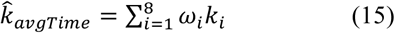

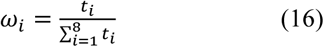

where *t*_*i*_ is the time spent interacting with the stiffness level *k*_*i*_. The participants were instructed to make discrete movements into the force field, which had required depth and time ranges that needed to be satisfied for the probe to be considered successful. Hence, occasionally there were trials in which a participant interacted more than once with the same stiffness level before achieving a successful probing movement. The participants likely updated their perceptual estimation about the stiffness during these ‘unsuccessful’ probing movements. The entire duration of the interaction with a given stiffness level was taken into account in this model.

**Average** – the average of the four stiffness levels:

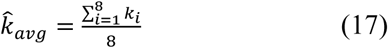

**Serially Weighted Average** – based on [11], in trials in which the *comparison* force field preceded the *standard*, unequal weights were attributed to the different stiffness levels, with the highest weight given to the first stiffness level and the lowest to the last stiffness level. On the other hand, if the *standard* was presented first, all eight stiffness levels were given equal weighting.

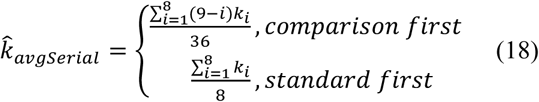

We then compared the stiffness value that was computed for a given model, 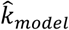, to the stiffness level of the *comparison* force field in that trial, and checked which value was higher.

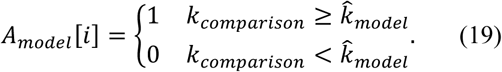

To calculate the score of the model, we compared the model’s answer to that of the participant. If the two were the same, the model received a point.

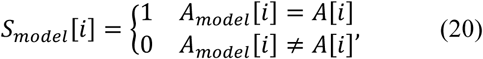

where *A*[*i*] is a participant’s response for a given trial, as defined in Equation (4). We analyzed the models’ predictions for each of the trials and calculated the number of trials in which the model corresponded with a participant’s response. The final step was normalizing the number representing the amount of corresponding trials to a score out of 100. We did so by dividing the final value by the total number of trials, N, and multiplying the result by 100.

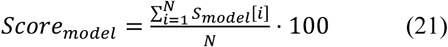

We performed this analysis for each of the participants and calculated the average score of the model.

To assess the statistical significance of the differences between the model scores, we used a repeated-measures General Linear Model with the MATLAB statistic toolbox (2018a), in which the independent variables were the model number (categorical, df=8), and the participants (random, df=9). The dependent variable was the model score, and we compared between every two models using post hoc t-tests (a total of 36 t-tests) with the Holm-Bonferroni correction method for multiple comparisons. We present the p-values after this correction (*p*_*corrected*)_, and therefore the threshold significance level following the correction is 0.05.

#### 3. Grip Force Control

We recorded the grip force (GF) data and filtered it using the MATLAB function *filtfilt* with a 2^nd^ order Butterworth low-pass filter, with a cutoff frequency of 15Hz, resulting in a 4^th^ order filter, with a cutoff frequency of 12.03 Hz. We separated the trials into four groups according to the different variability levels. We examined the GF applied by the participants and the trajectories of the vertical position of the hand (y axis) for every trial in each of the variability conditions. Classic analyses in the study of grip force control are: (i) to examine the grip force in relation to the load force by analyzing the ratio between the peak GF and peak load force (LF) [10, 27, 28]:

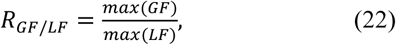

and (ii) to fit a linear regression model in the grip force – load force plane [22, 27, 28] :

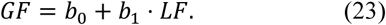

However, we will now show that varying the stiffness level between probing movements, as in our study, necessitates slightly modifying the approach; we analyzed the GF with respect to the vertical position of the hand rather than relative to the LF. We will first explain why the classical analysis is less appropriate in our study, and then present the rationale of the modified analysis.

The regression in (23) is fit under the assumption that participants estimate the anticipated load force, 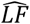 based on previous interactions and plan their grip force according to:

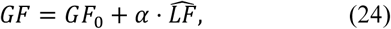

where *GF*_0_ is the grip force baseline and *α* is a coefficient which is affected by the friction at the contact interface as well as a safety margin. The safety margin is determined both by the participants’ uncertainty regarding the load force dynamics and the contact interface properties. As the participants interacted with an elastic force field, the assumption is that the anticipated load force is estimated according to:

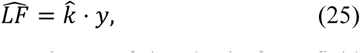

where 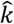 is the participants’ estimate of the elastic force field stiffness, and *y* is the planned penetration distance. Substituting (25) into (24) yields:

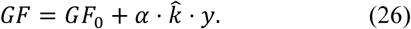

The actual load force can be described as *LF* = *k*. *y*, where *k* is the actual force field stiffness. Therefore: we rewrite (26) as:

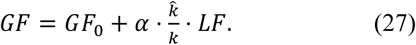

Comparing (23) and (27) highlights that the intercept of the regression in (23) represents a grip force baseline that is unaffected by the planned interaction with the elastic force field. On the other hand, the slope, which represents the grip force modulation in anticipation of the load force, is affected by the estimates of both the elastic force field stiffness and the contact dynamics.

Commonly, *k* in (27) is deterministic [27, 28, 46]. However, in our case, *k*∼*N*(85, *σ*), meaning k is not deterministic. In our analysis, we wish to quantify the effect of the variability on the measure of uncertainty experienced when estimating 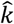. If we were to use the classical analysis, as in (23), in addition to the effect of the estimated stiffness value, the slope would also be affected by the variability in *k* (as demonstrated in (27)). However, if instead we compute a regression of the grip force as a function of the vertical position of the hand (*y*), as in (26), the slope will be affected solely by the estimates of the elastic force field stiffness levels and the contact dynamics. Hence, analyzing the grip force in relation to the vertical position results in a coefficient that is analogous in principle but not in units to the LF coefficient obtained in classic studies (as in equation (27) in the event of a deterministic k). When performing the regression as a function of the LF, the slope has no units ([au]), whereas calculating the regression as a function of the position leads to units of [N/m] for the slope. Nonetheless, comparing between the different variability levels’ regression slopes obtained in our analysis reflects the effect of the different variability levels on the modulation component of the grip force. Similar reasoning can be devised for the ratio between the peak GF and peak LF in (22), such that the appropriate analysis in our study is:

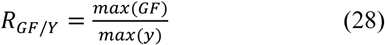

A limitation which arises when using this new approach is that it does not allow for comparing the slope and ratio values obtained using our analyses to those in the literature.

To gain insight into the changes that occurred in the grip force throughout the complete interaction with the *standard* force field, we separated between the different probing movements, and compared the grip force applied in the first probing movement to the grip force applied in the last probing movement. We identified the start and end of each probing movement by analyzing the trajectories of the vertical position of the hand. A probing movement began when participants crossed the boundary of the elastic force field (i.e., entered the negative half of the vertical axis) and ended when they exited the force field.

To quantify the effect of the variability condition on the grip force during the repeated exposure to the elastic force field, for the first and last probing movement into the *standard* force field in each trial we performed the two analyses described above: (i) We calculated the peak grip force-peak position ratio (*R*_*GF*/*Y*_, equation (28)) in the first and last probing movements in each of the test trials, and averaged these ratios for each of the variability conditions. The peak grip force-peak position ratio is a measure of the overall grip force, which is comprised of a baseline component and a modulation component. (ii) To separate the overall grip force into its two components, we fit a two degrees-of-freedom regression (intercept and slope) to the trajectory in the grip force-position plane (equation (26)).

To test the significance of the changes in the peak grip force-peak position ratio and in the intercept and slope of the grip force-position regression due to the variability condition and between the first and last probing movements, we fit a repeated-measures General Linear Model to each of these dependent variables separately using the MATLAB statistic toolbox (2018a). The independent variables were the variability condition (categorical, df=3), the probing movement (categorical, first or last, df =1), the interaction between them (categorical, df=3) and the participants (random, df = 9).

## III. Results

This experiment was completed over two sessions; half the participants completed the zero and medium variability trials first, while the other half did the low and high variability trials in the first session. To ensure that there was no effect of the order on the results, we compared between the results of the two groups and found no significant differences between them, both in the results of the perception analyses (rm-General Linear Model, PSE: main effect of ‘group number’: *F*_(1,8)_ = 0.03, *p* = 0.8722; JND: main effect of ‘group number’: *F*_(1,8)_ = 1.85, *p* = 0.1910), and in the results of the grip force analyses (rm-General Linear Model, Peaks Ratio: main effect of ‘group number’: *F*_(1,8)_ = 0.01, *p* = 0.9444; Intercept: main effect of ‘group number’: *F*_(1,8)_ = 0.27, *p* = 0.6169; Slope: main effect of ‘group number’: *F*_(1,8)_ = 0.10, *p* = 0.7625). Therefore, we combined the two groups for the remaining analyses in this paper.

### A. Perception and Uncertainty

In order to study the effect of the different levels of variability on the perceived stiffness, we fit psychometric curves to the responses of each of the participants in each variability level. The psychometric curves of a typical participant are presented in Fig. 3(a). The yellow psychometric curve represents the trials with zero variability (*σ* = 0. *JND*_*L*_ = 0 *N*/*m*), and the different shades of green show the results of the non-zero levels of variability [low (*σ* = 1. *JND*_*L*_ = 13), medium (*σ* = 2. *JND*_*L*_ = 26 *N*/*m*) and high (*σ* = 3. *JND*_*L*_ = 39 *N*/*m*)]. Using the psychometric curves, we calculated the Points of Subjective Equality (PSE) and the Just Noticeable Differences (JND). An examination of the psychometric curves in Fig. 3(a) revealed no horizontal shifts of the green psychometric curves relative to the yellow curve. This indicates that the different levels of variability did not affect the perceived stiffness. The slope of the psychometric curve for the trials with the medium level of variability was less steep than those in the zero, low, and high variability levels. This decrease indicates an increase in the JND, or the uncertainty.

**Fig. 3.**
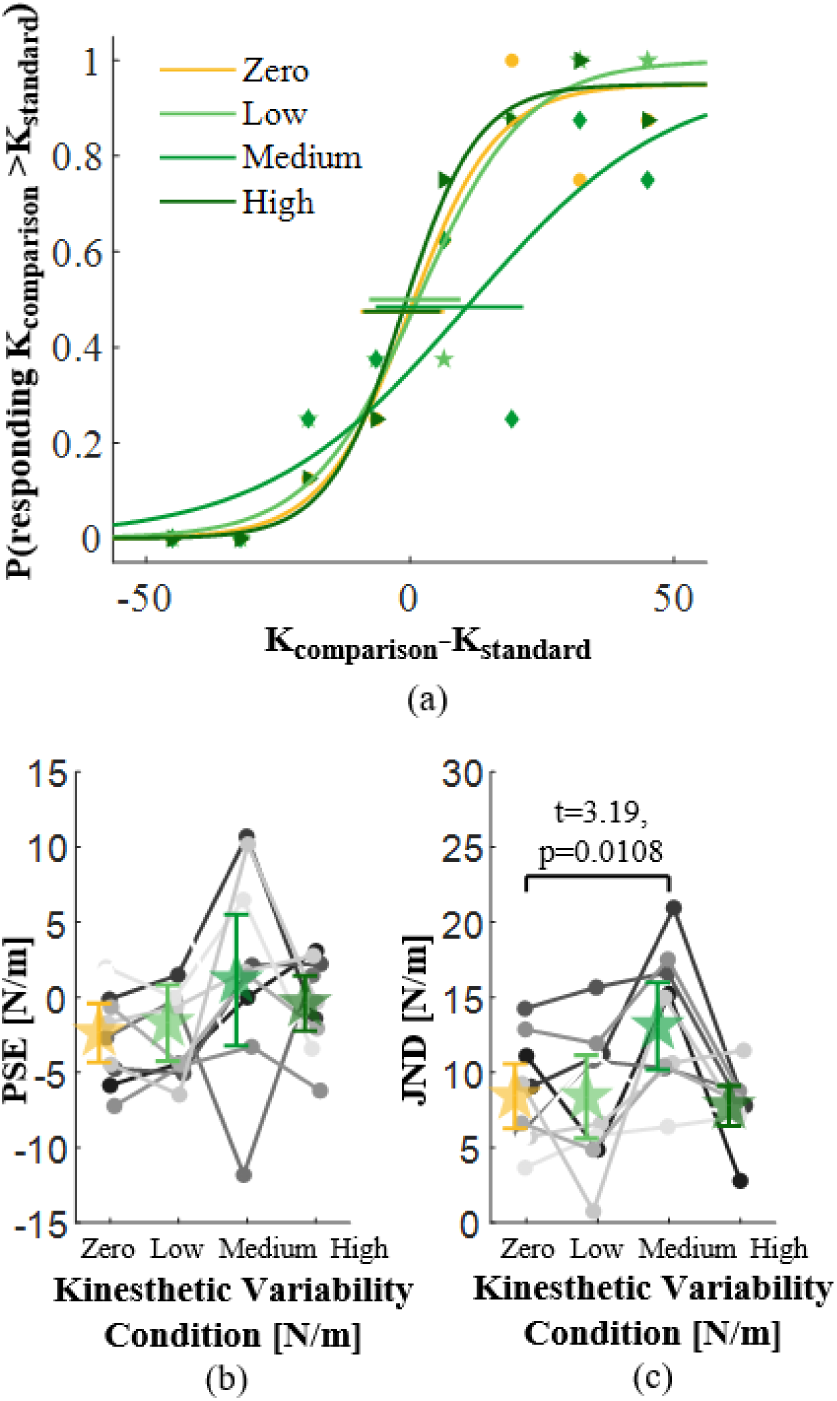
The effect of between-probe variability on the perception of stiffness. The yellow lines and symbols represent the trials with no variability and the three shades of green are for the three variability conditions – low, medium, and high. (a) The psychometric curves of a typical participant. The horizontal lines show the 95% confidence intervals for the PSE values. (b) The PSE (c) and JND values as a function of the different variability levels. The circles and lines with different shades of gray represent the data of each of the participants (N=10), and the colored stars and error bars show the average values across all the participants and the 95% confidence intervals, respectively.

The results of all the participants [Fig. 3(b) and 3(c)] were similar to those of the typical participant. Fig. 3(b) presents the PSE values of each of the participants, and their means, in the four variability conditions. We found no difference between the PSE values for the different variability conditions (PSE, rm-General Linear Model, main effect of ‘variability condition’: *F*_(3,27)_ = 1.1408, *p* = 0.3504), meaning the between-probe variability did not increase or decrease the perceived stiffness. Fig. 3(c) depicts the JND values of each of the participants in the four variability conditions, and their means. We observed a significant effect of the variability level on the JND (JND, rm-General Linear Model, main effect of ‘variability condition’: *F*_(3,27)_ = 5.6900, *p* = 0.0037). The three planned t-tests revealed an increase in the JND in trials with the medium level of variability relative to the trials with no variability (*t*_27_ = 3.19, *p*_*corrected*_ = 0.0108). Therefore, we conclude that the medium level of between-probe variability reduced the participants’ discrimination accuracy and caused them to be less precise in their answers.

### B. Models for Serial Stiffness Levels Integration

While the increased JND that we observed could stem from uncertainty caused by the variability in the stiffness level within a trial, a similar increase would be observed if the participants ignored some of the probing movements, and took only some of the stiffness levels into account when assessing the *standard* stiffness (for instance, only the first or last stiffness level). In this case the increase in the JND would be due to the perceived stiffness being distributed normally with the respective standard deviation across trials instead of within each trial. Although this would lead to a similar increase in the JND, it would not reflect an increase in the uncertainty due to the between-probe variability within the trial. The next step was therefore to ascertain that participants used all, or at least most of, the different *standard* stiffness levels presented in each trial when assessing the *standard* stiffness level. We did so by creating different models for several potential strategies participants may use.

Fig. 4(a) presents the scores of the different models, where the x-axis represents the model number. The model names and numbers are presented in Table 1, along with the mean score of each model and 95% confidence interval. The highest scoring models were the three models which averaged all eight stiffness levels, *Serially Weighted Average, Average*, and *Time Weighted Average*. The difference between the scores of the various models was statistically significant (scores, rm-General Linear Model, main effect of ‘model’: *F*_(8,72)_ = 84.62, *p* < 0.0001).

**TABLE 1.**
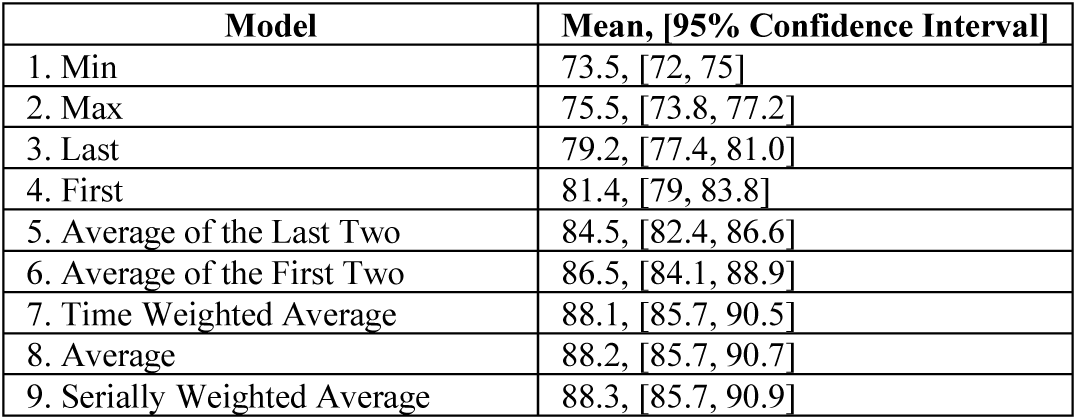
MODELS AND THEIR SCORES.

**Fig. 4.**
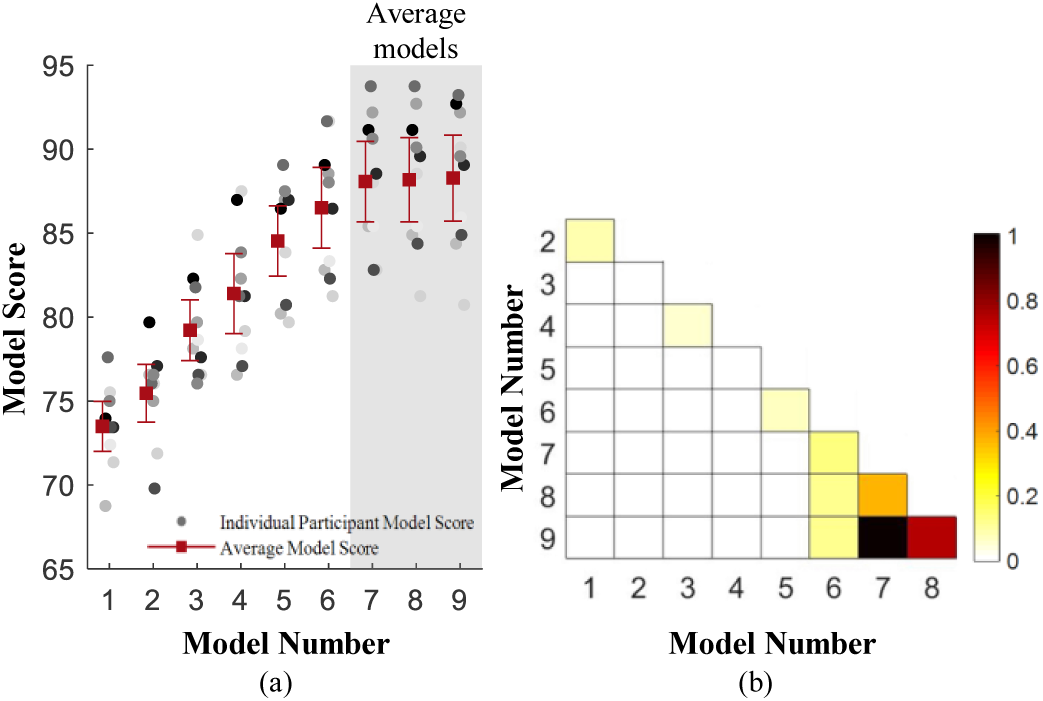
Computational models for serial stiffness levels integration. (a) The model scores for each participant and the mean scores across participants. The gray circles present the model score of each of the participants. The red squares show the mean score of each of the models and the error bars are the 95% confidence intervals. The ordinate is the score out of 100. The abscissa is the model number, in accordance with Table 1. The shaded gray region emphasized the three models according to which an average of all the stiffness levels was computed. (b) The results of post hoc t-tests, with Holm-Bonferrni corrections, between each two models. The different colors represent the corrected p values, where a paler color represents a lower p value, and a darker color signifies a higher p value (corresponding to the colorbar on the right).

To compare between each two models, we performed post-hoc t-tests, using the Holm-Bonferroni correction method for multiple comparisons, and present the corrected p values in the heatmap in Fig 4(b). A paler color (i.e., more similar to white) represents a lower p value, whereas a darker color signifies a higher p value. We found that the *Serially Weighted Average, Average*, and *Time Weighted Average* models scored significantly higher than all the other models, except for the Average of the First Two model, whose score was lower but not statistically significantly. These results indicate that participants likely took into account all the stiffness levels presented by the *standard* object when assessing its stiffness, leading us to believe that the observed increased uncertainty is the result of the between-probe variability.

### C. Grip Force Control

Fig. 5(a) presents examples of position and grip force trajectories of a typical participant during a typical trial (both *standard* and *comparison* force fields) in the zero variability condition. These trajectories clearly show that participants maintained a grip force-position modulation during the interaction with the force fields. Furthermore, consistently with prior literature [25-27], the grip force modulation began before the initial contact with the elastic force field, suggesting that it was at least partially predictive.

**Fig. 5.**
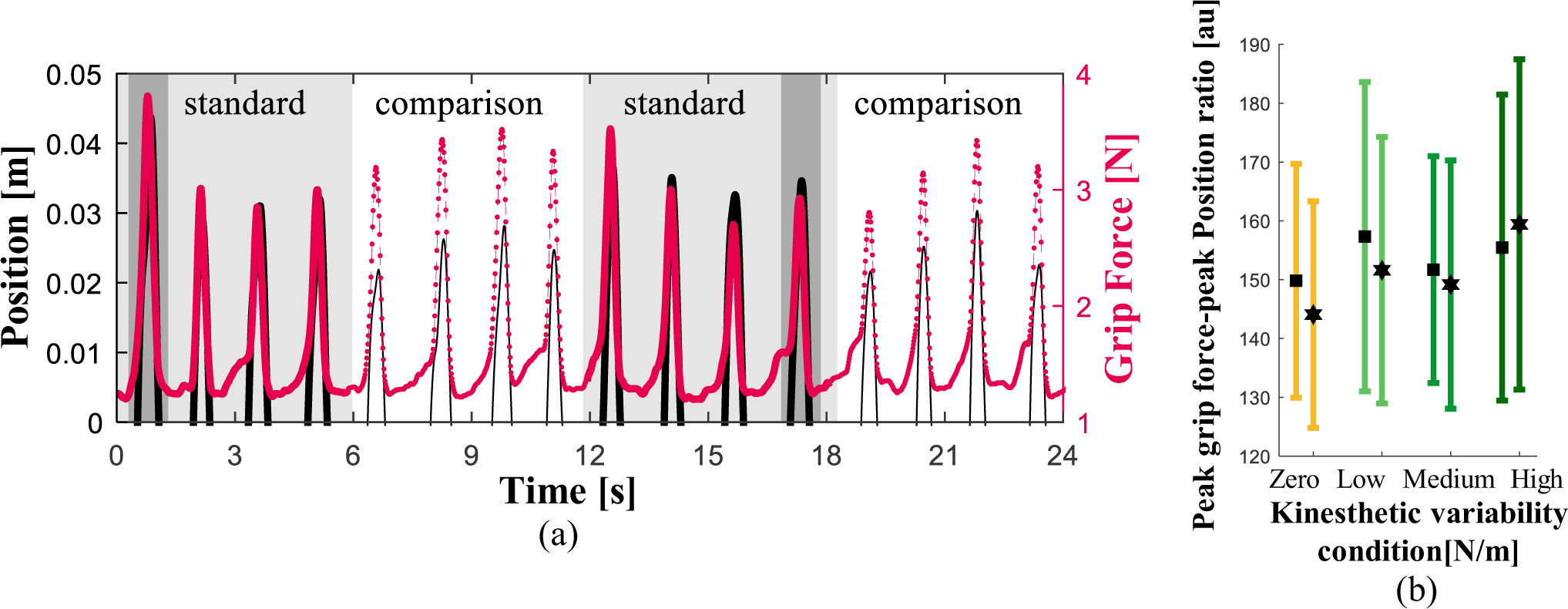
Analysis of grip force trajectories with respect to position trajectories. (a) Example of position (black lines) and grip force (red lines) trajectories of a typical participant during a zero variability trial. The light gray shaded regions highlight the probing movements into the standard force field,. The dark gray shaded regions highlight the first and the last probing movements, for which we analyzed the peak grip force-peak position ratio. (b) Analysis of peak grip force-peak position ratio for the first and last probing movements, averaged across all the participants for the four variability conditions. The vertical lines show the 95% confidence intervals.

In order to quantify the effect of the different variability levels on the grip force control, we analyzed the peak grip force-peak position ratio [Fig. 5(b)[. To separate the grip force into its two components (baseline and modulation), we performed a regression of the grip force as a function of position, and computed the intercept and slope in each variability condition [Fig. 6]. To compare between the effects of the different variability levels on early and late interactions, we performed these analyses for the first and last probing movements.

**Fig. 6.**
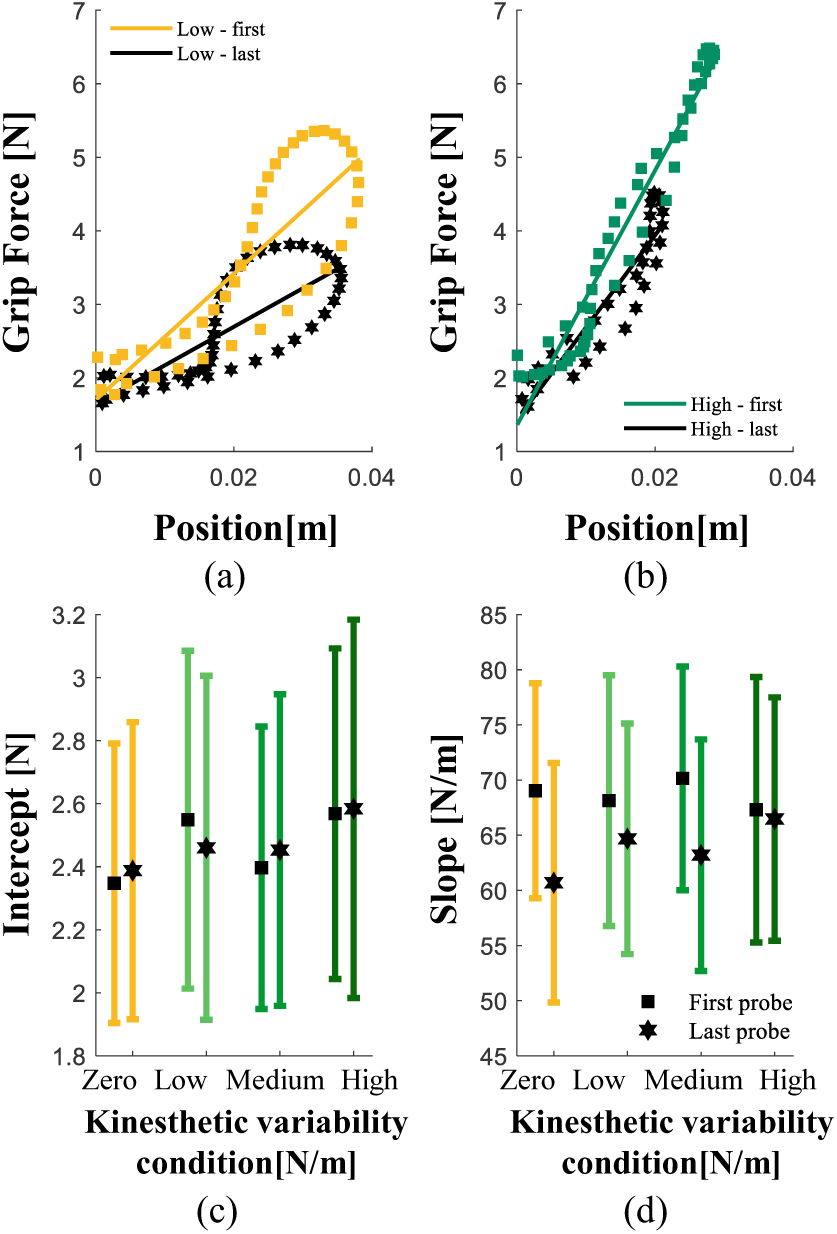
Grip Force-Position Regression analysis. (a-b) Examples of grip force-position trajectories and fitted regression lines from the first (colored) and last (black) probing movements from in zero and high variability trials, respectively. (c) Intercept and (d) slope of the regression lines that were fitted to the grip force position trajectories from the first and last probing movements in each trial, averaged across all the participants for the four variability conditions. The vertical lines show the 95% confidence intervals.

We observed a slight decrease in the peak grip force-peak position ratio for the zero, low and medium variability conditions, and a small increase for the high variability condition (Fig. 5(b)). However, these differences were not statistically significant (rm-General Linear Model, main effect of ‘variability condition’: *F*_(3,63)_ = 1.44, *p* = 0.2397; main effect of ‘probing movement’: *F*_(1,63)_ = 0.43, *p* = 0.5123; interaction between ‘variability condition’ and ‘probing movement’: *F*_(3,63)_ = 0.35, *p* = 0.7857).

Fig. 6(a) and 6(b) show examples of a typical participant’s trajectories in the grip force-position plane for the first and last probing movements in a zero and a high variability trial, respectively. These figures show no difference between the intercept of the first and last probing movements in both of the displayed variability conditions. When examining the trajectory slopes, we noted a decrease between the first and last probing movements. Furthermore, a comparison between Fig. 6(a) and 6(b) revealed a larger decrease in the slope of the zero variability condition relative to that of the high variability condition.

To assess the results of all the participants, we separated the trials according to the variability condition and, in each condition, averaged all the participants’ intercept and slope values from the first and last probing movements in all the trials. Fig. 6(c) displays the intercept component; similar to the results seen in the individual trajectories [Fig. 6(a) and (b)], we observed no effect of the variability condition and the probing movement (rm-General Linear Model, main effect of ‘variability condition’: *F*_(3,63)_ = 1.01, *p* = 0.3947; main effect of ‘probing movement’: *F*_(1,63)_ = 0, *p* = 0.9515; interaction between ‘variability condition’ and ‘probing movement’: *F*_(3,63)_ = 0.13, *p* = 0.9425) on the intercept. The lack of difference between the intercept in the first and last probing movements in the zero variability condition was an unexpected result, as we had anticipated that participants would decrease their grip force baseline with repeated interactions with the force field [28].

An analysis of the slope component in Fig. 6(d) showed a decrease between the first and the last probing movements (rm-General Linear Model, main effect of ‘probing movement’: *F*_(1,63)_ = 5.51, *p* = 0.022). This decrease indicates a reduction in the contribution of the modulation component to the control of grip force with the repeated interactions. Similar to the intercept analysis, we did not find a difference between the average slopes of the four variability conditions (rm-General Linear Model, main effect of ‘variability condition’: *F*_(3,63)_ = 0.19, *p* = 0.9004; interaction between ‘variability condition’ and ‘probing movement’: *F*_(3,63)_ = 0.66, *p* = 0.5785).

## IV. DISCUSSION

In this work we strived to design a method of introducing uncertainty into the haptic information without distorting (e.g., adding noise to) the haptic feedback. We hypothesized that varying the stiffness level of elastic force fields between consecutive interactions may create haptic uncertainty that would come across in both stiffness perception and grip force adjustment. If this were to be the case, it would enable the study of perception and action creation without necessitating changes in the quality of the haptic information supplied in each individual interaction.

Our results revealed that the medium level of between-probe haptic variability created uncertainty in the perception of stiffness, quantified by an increase in the JND. Furthermore, by testing different models for the integration of serial probing movements with varying stiffness levels, our analysis indicates that participants appeared to have taken all the different stiffness levels applied by a given force field into account when forming their perception. Our examination of the grip force applied by the participants showed no difference between the grip force applied in the different variability conditions (zero, low, medium and high). Moreover, we found no decrease in the grip force baseline with repeated exposure to the force field in all the variability conditions, whereas the modulation component did decrease. These results may be due to a global effect of the uncertainty on the control of grip force that was experienced in all the variability conditions.

Perceptual uncertainty was apparent in the event of the medium level of variability. This result demonstrates the efficacy of between-probe haptic variability as a method for creating uncertainty and indicates that the medium level of variability is appropriate for producing the desired uncertainty. This induced uncertainty is in accordance with previous studies which showed that the addition of noise to sensory inputs can impair the discrimination ability [12, 13, 47, 48]. Ernst and Banks [12] found that adding varying levels of noise to the visual input increased participants’ uncertainty regarding the visual information. A study by Machworth et al. [48] suggested that adding visual noise lowered participants’ ability to detect whether visual displays were identical or different. Dallman et al. [47] demonstrated a reduced precision in speed discrimination due to the addition of vibratory noise. Gurari et al. [13] showed that adding haptic white noise led to a degradation in participants’ ability to identify the magnitude of stiffness and force stimuli. While the discrimination sensitivity of the participants in our study was impaired by the addition of the variability, the perceived stiffness was not biased. That is, the addition of the variability did not lead to an increase or decrease in the perceived stiffness of the force field. This is consistent with the findings of Zanker et al. [49] who created different levels of noise in visual speed feedback. They observed that while the noise had no effect on the perceived speed, the noise impacted the certainty. The results of our study are consistent with those of previous works, despite the novel method we used to introduce uncertainty. Whereas prior research has altered sensory inputs by adding noise, we chose to introduce between-probe variability. This method allows for obtaining the desired uncertainty without distorting the information presented within each probing movement.

To confirm our conclusions about the creation of perceptual uncertainty, it was critical to ascertain that the participants used the information from at least most of the *standard* probing movements in a trial when choosing the stiffer force field. Had participants based their responses solely on a single stiffness value, e.g. the first or the last, a similar increase in the JND would be observed. This increase, however, would be the result of the variability of the single stiffness level which they took into account between trials, and not due to the within-trial variability. For example, if participants were to base their estimate only on the first stiffness level, as there are trials in which this stiffness level is below 85 N/m, and trials in which it is above, we would observe an increase in the JND that does not indicate uncertainty due to the within-trial variability.

With the goal of uncovering how serial information is combined when experiencing haptic variability, we created computational models for various potential strategies participants may use, and ran trial-by-trial simulations in which we evaluated the compatibility of each model with the participants’ responses. We found that the models which best predicted the participants’ choices were the models in which participants took all eight stiffness levels into account and averaged them. Specifically, the *Serial Weighted Average Model* scored slightly higher than the other two average models. This result is consistent with the results found in the work of Metzger et al. [11]. They found that the nervous system attributes decreasing weights to the different stiffness levels based on their serial placement in the probing sequence, with the highest weight given to the stiffness level experienced first and the lowest to the last. The superiority of the average models indicates that our results did not stem from participants negating part of the haptic information supplied by the *standard* force field and that the observed increase in the JND may be due to uncertainty caused by the within-trial variability.

Unlike the medium variability, the low and high levels of variability did not create haptic uncertainty. The lack of effect of the low variability level may be due to insufficient differences between the stiffness levels presented in the consecutive probing movements. Indeed, as the stiffness levels in this condition were selected from a normal distribution whose standard deviation was 1·JND_L_ [44], the difference between most of the stiffness levels applied by a given force field was within the range in which participants may have been unable to distinguish between them. The fact that the high level of variability did not create uncertainty was an unexpected result — we hypothesized that the high level would increase the uncertainty more than the medium variability. The reason behind this result remains an open question and necessitates further investigation.

While the perceptual effect was evident only in the medium variability condition, uncertainty in the grip force may have been present in all four variability conditions. A similar peak grip force-peak position ratio was observed in all of the variability conditions, implying that participants may have experienced general uncertainty due to the variability and therefore maintained a relatively constant safety margin throughout the entire experiment. This may be due to participants being uncertain about what level of stiffness to expect in each subsequent probing movement

To dissociate the effect of the variability on each of the two grip force components, we individually examined the baseline and modulation. We found that participants applied a similar grip force baseline in all four variability conditions. This result is in contrast to those of Hadjiosif and Smith [42], who showed an increase in the grip force baseline due to variability in velocity-dependent load forces. An examination of the modulation component presented a similar picture of equal values across the different variability conditions. These results correspond with the result of the peaks ratio analysis; that is, participants appeared to maintain a safety margin due to general uncertainty.

We investigated the progression of the grip force baseline and modulation with repeated exposure to the force field. When comparing between the first and last probing movements, our analysis revealed no decrease in the grip force baseline with subsequent probing movements in all four variability conditions. This result was unexpected; Leib et al. [28, 35] and Farajian et al. [27] observed a decrease in the grip force baseline with repeated interactions with a given force field. They attributed this finding to an increase in the certainty regarding the load force which resulted in a reduction in the safety margin. We expected to find a decrease in the grip force baseline at least in the zero variability condition, as in this condition, the stiffness level remained constant. As we observed no decrease in the grip force baseline with repeated interactions with a given force field in all the variability conditions, we believe that there was a general effect of uncertainty on the control of grip force throughout the entire experiment, likely causing participants to preserve a high safety margin. This view is corroborated by our observation that the modulation component decreased significantly with repeated interactions in all four variability conditions. Thus, as participants continued to experience varying stiffness levels between consecutive probing movements, their grip force controller relied less on their estimations of the expected load force, and became more similar to the control of grip force in the absence of load force information [50]. To summarize, the uncertainty appeared to affect both the grip force baseline and the modulation in anticipation of the load force. The first manifested in a lack of reduction of the safety margin element of the grip force baseline with repeated interactions, and the latter, in a decrease of the modulation component.

## V. CONCLUSIONS

In this study, we conducted an experiment that allowed for the characterization of the effect of between-probe variability in the stiffness of elastic force fields that are rendered by a haptic device on stiffness perception and grip force control. Our findings indicate that perceptual uncertainty is created in the event of the medium variability level, whereas uncertainty in the grip force is evident in all four variability conditions. The latter is likely due to the distinct variability conditions causing a general effect of motor uncertainty that affected the control of grip force throughout the experiment rather than on a trial-by-trial basis. Furthermore, our perceived stiffness appears to be a weighted average of the different stiffness levels applied by a given force field. In addition to contributing to the understanding of the processing of haptic information, our study presents a method for creating haptic uncertainty while presenting undistorted haptic information.

